# Targeted single-cell RNA-seq identifies minority cell types of kidney distal nephron that regulate blood pressure and calcium balance

**DOI:** 10.1101/2020.07.19.209627

**Authors:** Lihe Chen, Chun-Lin Chou, Mark A. Knepper

## Abstract

A major objective in modern biology is generation of comprehensive atlases of various organs identifying all cell types and their expressed genes. In kidney, extensive data exists for proximal tubule and collecting duct cells, but not for non-abundant intermediate epithelial cell types. Here, we coupled a FACS-enrichment protocol with single-cell RNA-seq analysis to profile the transcriptomes of 9099 cells from the nephron region adjacent to the macula densa. Clusters containing *Slc12a3*^+^/*Pvalb*^+^ and *Slc12a3*^+^/*Pvalb*^-^ cells were identified as DCT1 and DCT2 cells. The DCT1 cells appear to be heterogeneously associated with variable expression of *Slc8a1, Calb1*, and *Ckb* among other mRNAs. No DCT2-specific transcripts were found. The analysis also identified two distinct cell types in the *Slc12a1*^+^ portion of Henle’s loop as well as *Nos1*^+^/*Avpr1a*^+^ macula densa cells. Thus, we identify unexpected cell diversity in the intermediate region of the nephron and create a web-based data resource for these cells.

## INTRODUCTION

The mammalian kidneys play a crucial role in regulation of body fluid composition and blood pressure. These functions are largely achieved by control of sodium reabsorption across the epithelia of renal tubules^1, 2^. There are at least 14 different renal tubule segments, each with characteristic cell types with distinct gene expression profiles and function^3, 4^. Of particular importance is the distal convoluted tubule (DCT), a short segment that mediates fine regulation of sodium ion transport^5^. The apical component of sodium transport across the DCT occurs via the Na^+^-Cl^-^ cotransporter NCC (Slc12a3), which is exclusively expressed in DCT cells and the chief target of thiazide diuretics^6^, a staple in the treatment of many forms of arterial hypertension. The DCT is believed to be heterogeneous and is separated into at least two subsegments, DCT1 and DCT2^7, 8, 9^. Both express Slc12a3. DCT1 but not DCT2 expresses the Ca^2+^-binding protein, parvalbumin (Pvalb). DCT2, in contrast to DCT1, expresses the epithelial sodium channel (ENaC)^9^. More recent observations indicate that the DCT2 also expresses the Ca^2+^ transporter, Trpv5^10^, and has been proposed to be a site of transepithelial Ca^2+^ transport^9, 11^. However, this DCT classification is largely defined based on immunocytochemical results. It remains unclear how many cell types are in DCT and whether DCT1 and DCT2 are distinct segments or represent two ends of a continuum.

NaCl transport across the DCT was originally thought to be affected in part by binding of circulating aldosterone to the mineralocorticoid receptor (MR) in DCT cells^12, 13^. Aldosterone-MR signaling requires an enzyme 11-β-hydroxysteroid dehydrogenase 2 (Hsd11b2) to deactivate the more abundant circulating cortisol, which has an equal binding affinity to MR. However, immunocytochemical and in situ hybridization methods to localize Hsd11b2 in DCT cells have been indicative of variable expression along the DCT with detectable levels only in the distal-most DCT^14, 15^. These findings raised doubts about a possible direct role of aldosterone in the early DCT, and whether the Hsd11b2 positive DCT corresponds to DCT2. Recent studies further suggest that Slc12a3 (NCC) abundance is regulated by aldosterone only indirectly via its propensity to cause hypokalemia^5, 16^. This process is thought to be controlled by multiple receptors, protein kinases (Wnk1, Wnk4, and SPAK), ubiquitin ligase components (Nedd4l, Klhl3, Cul3), potassium channels, and chloride channels^5, 16^. However, a systematic mapping of these elements in DCT is currently not available.

The progressive development and improvement of RNA-seq for comprehensive quantification of gene expression, especially small-sample RNA-seq in microdissected renal tubules^17^, and more recent single-cell RNA-seq techniques in kidney^18, 19, 20, 21, 22, 23, 24, 25^, have provided a deep analysis of gene expression in the kidney epithelial cells and have broadened our insights into renal cell identities and functions. The tubule microdissection method^26^, however, is unable to effectively isolate the DCT1 and DCT2 due to the overall shortness of DCT and the lack of a distinct transition point between DCT1 and DCT2. Despite great progress of comprehensive single-cell RNA-seq in the kidney (‘shotgun’ scRNA-seq), it devotes most of the sequencing reads to more abundant proximal tubule cells and non-epithelial cells, leading to analysis of only limited numbers of minority epithelial cell types. Thus, the heterogeneity of cell types like DCT cells as well as various cell types in the loop of Henle (for example, macula densa and cortical thick ascending limb cells) are not completely resolved. A solution to this dilemma is to enrich the target cells before analysis as previously introduced for scRNA-seq analysis of collecting duct cell types^19^.

Here, we describe an enrichment protocol prior to single-cell RNA-seq analysis and specially focused on the mouse nephron region spanning from the cortical thick ascending limb of Henle (CTAL) to the DCT. In parallel, we use small-sample RNA-seq to characterize the gene expression in all 14 renal tubules microdissected from mouse kidneys. We also provide web-based resources that allow users to explore and download data for future studies.

## RESULTS

To provide appropriate context for single-cell RNA-seq (scRNA-seq) analysis of cells in nephron region from cortical thick ascending limb of Henle (CTAL) to the distal convoluted tubule (DCT) to the connecting tubule (CNT) in mouse, we began by mapping transcriptomes in microdissected CTAL, DCT, and CNT tubule segments. We also carried out RNA-seq in all of the other segments including proximal tubule segments, thin limbs of Henle’s loop, and all collecting duct segments to provide a comprehensive transcriptomic landscape of mouse renal epithelial cells. To characterize the cellular composition and transcriptional profiles of the cells in the CTAL-DCT-CNT region, we developed a fluorescence-activated cell sorting (FACS) protocol and carried out single-cell RNA-seq analysis (scRNA-seq) of these cells (Fig. 1a). The resulting data are viewable at https://esbl.nhlbi.nih.gov/MRECA/Nephron/ and https://esbl.nhlbi.nih.gov/MRECA/DCT/.

**Fig. 1.**
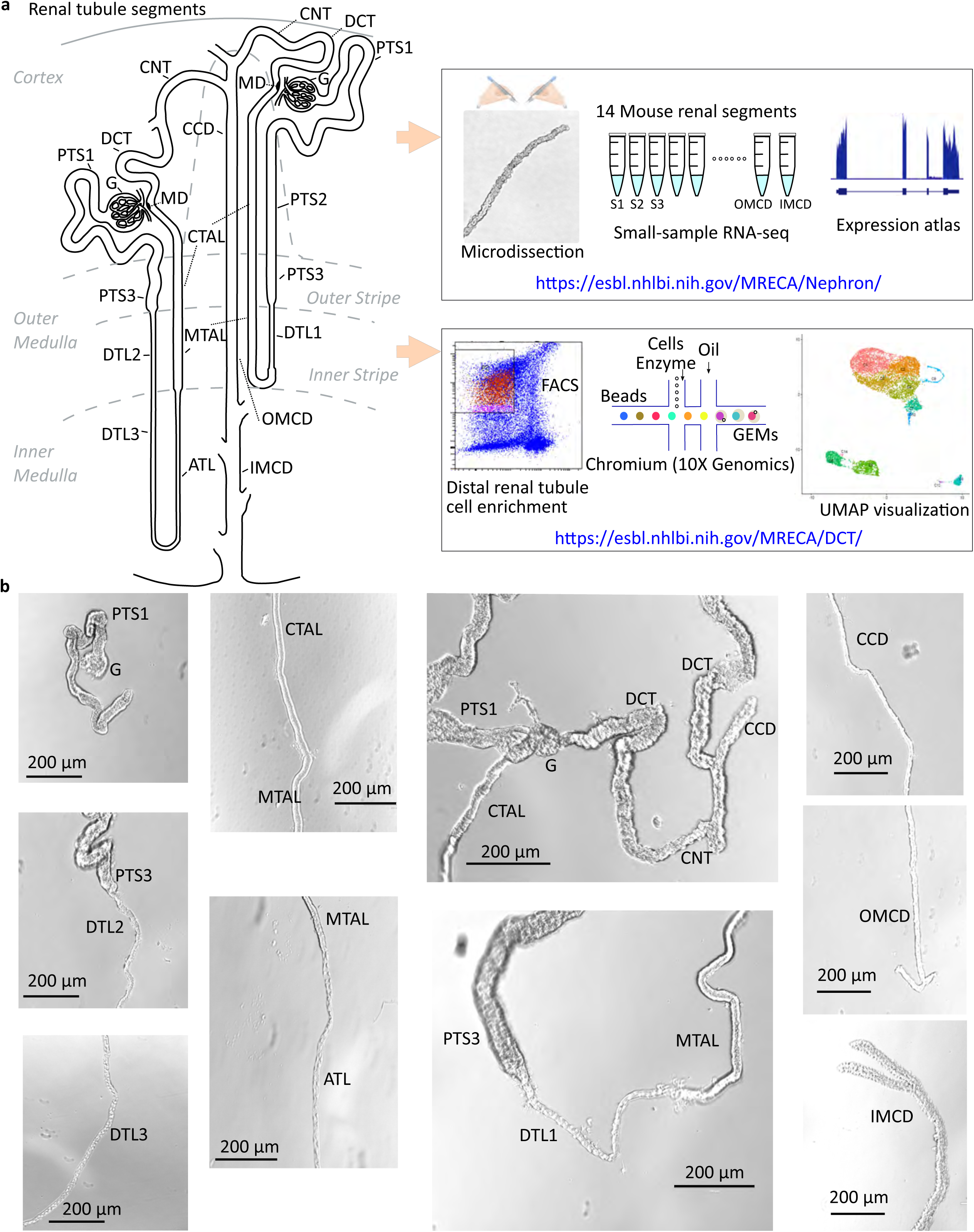
Overview of renal tubule cell nomenclature and experimental design. **a** The scheme^3, 4^ shows the connection of both a short-looped and a long-looped nephron to the collecting duct system. Small-sample RNA-seq coupled with microdissection was used to quantify gene expression in all 14 mouse renal tubule segments. An enrichment protocol was developed for single-cell RNA-seq analysis of distal convoluted tubule cells. The data are provided in user-friendly web sites. **b** Representative images of microdissected mouse renal tubule segments. The images were captured by the Invitrogen EVOS XL Core Cell Imaging System. Scale bars, 200 µm. G, Glomerulus; PTS1, the initial segment of the proximal tubule; PTS2, proximal straight tubule in cortical medullary rays; PTS3, last segment of the proximal straight tubule in the outer stripe of outer medulla; DTL1, the short descending limb of the loop of Henle; DTL2, long descending limb of the loop of Henle in the outer medulla; DTL3, long descending limb of the loop of Henle in the inner medulla; ATL, thin ascending limb of the loop of Henle; MTAL, medullary thick ascending limb of the loop of Henle; CTAL, cortical thick ascending limb of the loop of Henle; MD, mecula densa; DCT, distal convoluted tubule; CNT, connecting tubule; CCD, cortical collecting duct; OMCD, outer medullary collecting duct; IMCD, inner medullary collecting duct.

### Transcriptomic landscape of microdissected CTAL, DCT, and CNT

To characterize gene expression in CTAL, DCT, and CNT plus all other renal tubule segments, we performed renal tubule microdissection under a standard stereomicroscope (Fig. 1b) followed by small-sample RNA-seq (Supplementary Data 1). For DCT, it was impossible to see well-defined boundaries between DCT1 and DCT2, so we did not separately dissect these two. In parallel, we generated a separate transcriptome dataset for glomerulus (https://esbl.nhlbi.nih.gov/MRECA/G/). In total, we collected 93 samples from 53 mice (87 renal tubules plus 6 glomeruli) with an average of 85% uniquely mapped reads (Supplementary Fig. 1a) and an average of 3.7% mitochondrial fraction (Supplementary Fig. 1b), indicating high data quality.

As shown in Fig. 2a, the distribution of recognized kidney cell-type-specific markers along the renal tubule matched well with prior knowledge^27^. Specifically, the bumetanide-sensitive Na-K-2Cl cotransporter 2 (*Slc12a1*) was restricted to MTAL and CTAL, the thiazide-sensitive Na^+^-Cl^-^ cotransporter (*Slc12a3*) and the Ca^2+^-binding protein parvalbumin (*Pvalb*) were confined to DCT, while aquaporin-2 (*Aqp2*) was limited to CD segments including the CNT. Overall, these data illustrate the high degree of concordance among replicates, demonstrating the reproducibility and accuracy of the small-sample RNA-seq approach applied to microdissected tubules.

**Fig. 2.**
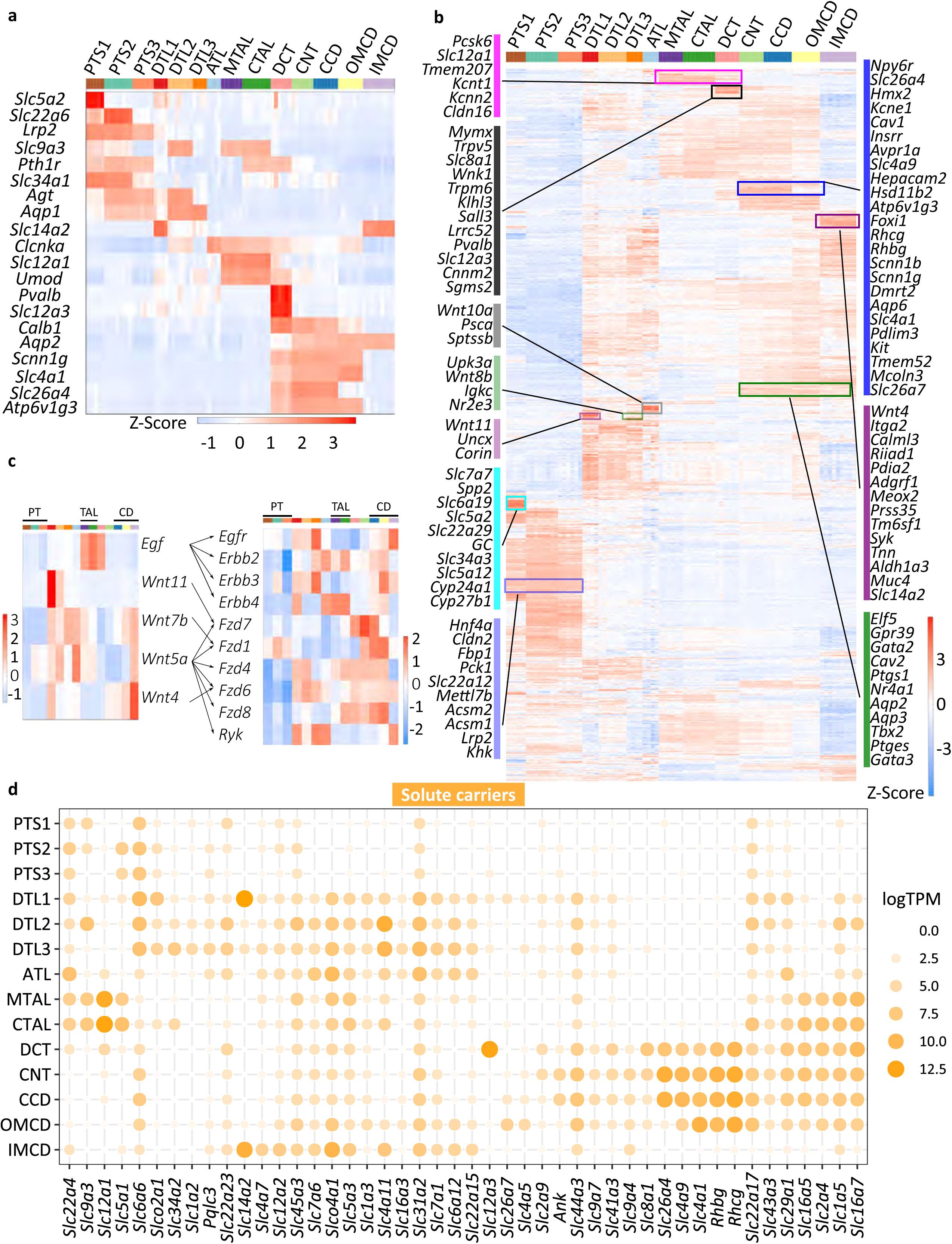
Gene expression pattern among mouse renal tubule segments. **a** Heatmap showing the expression pattern of classical renal tubule markers. Columns are individual renal tubule replicates and rows are marker genes. Color bar (top) indicates different renal tubule segments. Red color indicates high expression, and blue color indicates low expression. Z-Score was calculated from log_2_(TPM+1) using *scale* function. **b** Heatmap displaying the scaled expression pattern of 3709 DEGs along the renal tubule segment. The full gene list is provided in Supplementary Data 3. **c** Representative ligand-receptor expression patterns. Left heatmap shows the mean ligand expression; Right shows the receptor expression. The full list is provided in Supplementary Data 4. **d** Dot plot displaying major non-proximal tubule solute carriers along the mouse nephron. Dot size and color indicate expression [Log2(TPM+1)].

Next, we performed differential gene expression analysis on each renal tubule segment separately and identified 3709 differentially expressed genes (DEGs) (Supplementary Data 2). Based on the degree of similarity of their DEG expression patterns, three major renal tubule clusters were revealed (Supplementary Fig. 1c) in which the TALs, DCT, and CNT were grouped in one cluster, indicating a close segmental/spatial gene relationship between them^17, 28^. The transcripts accounting for these patterns are presented in Fig. 2b, many of which are products of novel segment-specific genes. The full list of these segment-specific genes is also provided in Supplementary Data 3.

In the CTAL-DCT-CNT region, some transcripts with less well-characterized function were selectively enriched. For example, TALs selectively expressed *Tmem207* and *Pcsk6* (proprotein convertase subtilisin/kexin-6). *Tmem207*, which codes for a single-pass type I membrane protein, showed restricted expression in kidney versus all other tissues (Genotype-Tissue Expression project, https://www.gtexportal.org/home/gene/TMEM207), suggesting a specialized role in the kidney. Pcsk6 acts as a secreted enzyme that activates atrial peptide converting enzyme, i.e. Corin, a serine peptidase, in heart, thereby regulating blood pressure^29^. Interestingly, very, very high levels of *Corin* were found exclusively in DTL1, where it can potentially be released by Pcsk6 derived from the neighboring MTAL. In DCT, some transcripts with no prior known functional roles in the DCT were identified. For example, LRRC52 is a K^+^ channel auxiliary gamma subunit that modulates the function of Slowpoke-homolog potassium channels including Maxi-K (Kcnma1)^30^ and Kcnu1^31^.

To further identify unique patterns of gene distribution from Fig. 2b, we mapped ligand/receptor pairs (Fig. 2c), solute carriers (Fig. 2d), transcription factors (TF) (Fig. 3a), and G protein-coupled receptors (GPCRs) (Fig. 3b) along the renal tubule and specifically focused on the CTAL-DCT-CNT region. Fig. 2c shows that *Egf* was predominately localized in TAL and DCT^32^, while its receptors were seen in various segments. This analysis also showed various Wnt protein transcripts among the thin limb segments. Other ligand-receptor pairs are listed in Supplementary Data 4. The major non-proximal selective solute carriers (transporters) are shown in Fig. 2d. Many of them match the current understanding of their localization and function. Examples include Na^+^/H^+^ exchanger 3 (*Slc9a3*), the bumetanide-sensitive Na-K-2Cl cotransporter (*Slc12a1*), and sodium/glucose cotransporter 1 (*Slc5a1*) in TAL; *Slc12a3* in DCT; and the Na^+^-Ca^2+^ exchanger (*Slc8a1*) *and* Glut9 (*Slc2a9*)^33^ in both DCT and CNT. Fig. 3a summarizes the distribution of major transcription factors along the renal tubule. Of particular interest are the transcription factors expressed in DCT. *Sall3*, a zinc-finger TF, was selectively expressed in DCT. *Emx1*, a homeobox TF, was also enriched DCT. It has been previously shown to be important for distal segment development in zebrafish embryonic kidney^34^. Several G protein-coupled receptors were strongly expressed in CTAL-CNT-DCT (Fig. 3b). Examples include parathyroid hormone 1 receptor (*Pth1r*)^35^ in PT, TAL and DCT; Prostaglandin EP_3_ receptor (*Ptger3*)^36^, calcium-sensing receptor (*Casr*)^37^, glucagon receptor (*Gcgr*)^38^, and arginine vasopressin receptor 2 (*Avpr2*)^39^ in TAL-DCT-CNT region. Immunofluorescence, coupled with in situ hybridization with RNAscope, confirmed the expression of *Ptger3* in some, but not all cells of the TAL (Fig. 3c) (see also below), supporting the fidelity of the data. The analysis also identified some GPCRs without well-characterized kidney functions including cholecystokinin A receptor (*Cckar*) in PT. A recent study indicated that it might regulate proximal tubular reabsorption^40^. Additional analyses were provided in this study including the distribution of aquaporins (Supplementary Fig. 1d) and ion channels (Supplementary Fig. 1e).

**Fig. 3.**
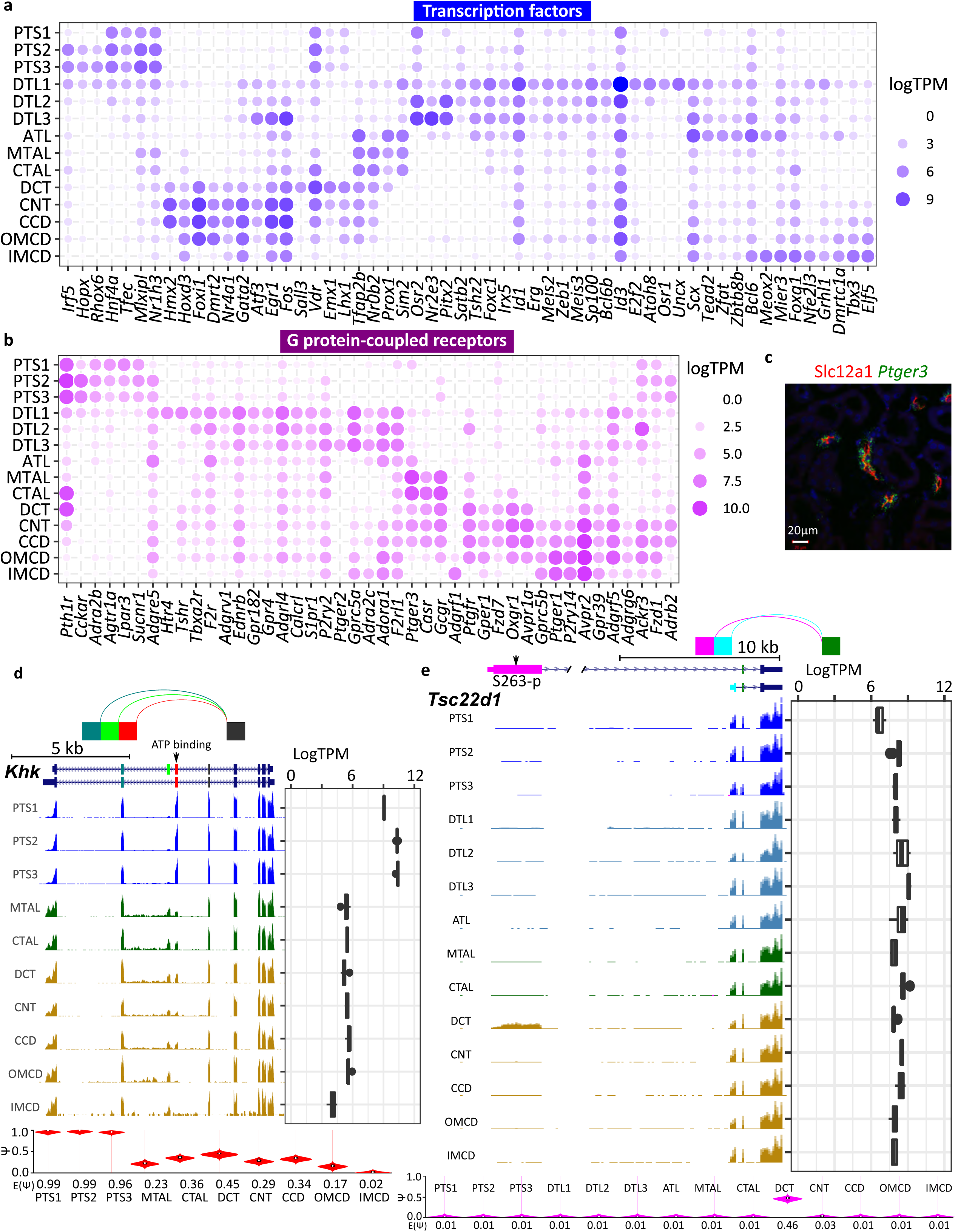
Gene expression pattern and alternative splicing. **a** Dot plot displaying important transcription factors along the mouse nephron segments. Dot size and color indicate expression [Log_2_(TPM+1)]. **b** Dot plot displaying important G protein-coupled receptors along the mouse renal tubule. Dot size and color indicate expression [Log_2_(TPM+1)]. **c** Immunostaining coupled with RNAscope showing *Ptger3* expression (green dots) in Slc12a1 cells (red) in mouse kidney sections. Scale bar, 20 µm. **d** Distribution of Khk isoforms. Mapping is visualized in the UCSC Genome Browser. The splicing graph at the top of the panel was generated using MAJIQ-SPEL. The paths on the splicing graph indicate the splicing junctions. The same color codes were used for exons in the splice graph and UCSC Genes Track. ATP binding site is indicated by the arrow. Khk gene expression is shown on the right. Quantifications of relative abundance (PSI) of red exon are shown on the bottom [sum of PSIs (dark green + green + red) =1]. PSI was calculated by MAJIQ and visualized by MAJIQ-SPEL. UCSC Genes Track scale, 5 kb. Track height scale, auto. **e** A similar analysis as in **d** was performed for Tsc22d1. Tsc22d1 shows a unique splicing isoform in DCT. Pink exon was mainly seen in DCT. UCSC Genes Track scale, 10 kb. Track height scale, auto. Quantifications of relative abundance (PSI) of purple exon are shown on the bottom [sum of PSIs (purple + cyan) =1].

In summary, we generated a comprehensive mouse renal tubule epithelial cell expression atlas (https://esbl.nhlbi.nih.gov/MRECA/Nephron/), which includes deep transcriptomes for the region of interest in this paper (CTAL-DCT-CNT) as well as those of all other tubule segments.

### The landscape of alternative splicing in mouse renal tubules

Our full-length RNA-seq protocol enables us to systematically map alternative exon usage along the renal tubule. We examined alternative splicing events using MAJIQ^41^, which quantifies reads that span splice sites (See *Method*). In total, we detected 931 confident splicing sites across 683 genes (Supplementary Data 5). Our data revealed distinct segment-specific isoforms of many genes including known isoforms of *Cldn10, Kcnj1* (ROMK), *Slc12a1* (NKCC2), *Wnk1, Stk39* (SPAK), and *Slc14a2* (Urea transporter A). Some novel isoforms were also identified in this analysis. Examples include ketohexokinase (*Khk*), which is abundantly expressed in proximal tubules and is critical for fructose metabolism, which was seen to have a dominant alternative isoform in proximal tubules (Fig. 3d). Specifically, the ATP binding domain-containing isoform needed for kinase activity was dominant in PTS1, S2, and S3, consistent with the known metabolic role of Khk in proximal tubules. We also identified a regulatory protein, *Tsc22d1*, showing a DCT-specific isoform with an extra 5’ coding exon (Fig. 3e). This DCT-specific isoform (ENSMUST00000048371, 1077 aa) is much longer (by 934 amino acids) than the isoform seen in the other tubule segments (ENSMUST00000022587, 143 aa). This short isoform is believed to function as a leucine zipper transcription factor. The extended N-terminal region in the DCT-specific isoform shows many low complexity regions (typically associated with RNA binding) and contains a Pat1 domain implicated in regulation of RNA stability (see UniProt record E9QLZ1). This observation suggests specialized roles of Tsc22d1 in DCT related to post-transcriptional regulation of gene expression. Other alternative splicing examples (For example, *Tpm1* and *Pth1r*) were provided in Supplementary Fig. 2. Collectively, these data provide a novel resource available in Supplementary Data 5 and at https://esbl.nhlbi.nih.gov/MRECA/Nephron/ describing variations in exon usage along the renal tubule.

### Single-cell RNA-seq of distal convoluted tubule (DCT) cells

DCT is generally subdivided into DCT1 and DCT2^7, 8, 9^. Both express Slc12a3. DCT1 but not DCT2 expresses Pvalb (Fig. 4a) while DCT2 selectively expresses ENaC (see below). The fairly short nature of the DCT2 and ill-defined DCT1/DCT2 and DCT2/CNT boundaries (Fig. 1b) prevented us from identifying a DCT2 transcriptome in microdissected tubules. Thus, we sought to use a FACS approach for the enrichment of DCT cells for scRNA-seq (i.e. Slc12a3^+^ cells), based on a strategy similar to one successfully applied to study heterogeneity of CD cells^19^. We began by dissociating mouse renal cortical tissue into single-cell suspensions using a cold-active protease^18^. The FACS sorting used positive selection with an antibody to Embigin (*Emb*), which encodes a single-pass type I membrane protein present in all cortical renal tubules except for proximal tubules (Fig. 4b), thus largely eliminating abundant PT cells^27^, while enriching DCT cells. We also used lectins for negative selection, namely LTL (Lotus tetragonolobus lectin) and PNA (peanut lectin), to remove remaining proximal and collecting duct cells (Fig. 4c). scRNA-seq utilized the 10X Chromium platform (Fig. 4d). We carried out 3 independent runs with 3000-6000 cells each as input. After data quality control and batch effects removal (Supplementary Fig. 3a) (See *Method*), we detected 17655 genes across 9099 cells (Supplementary Fig. 3b).

**Fig. 4.**
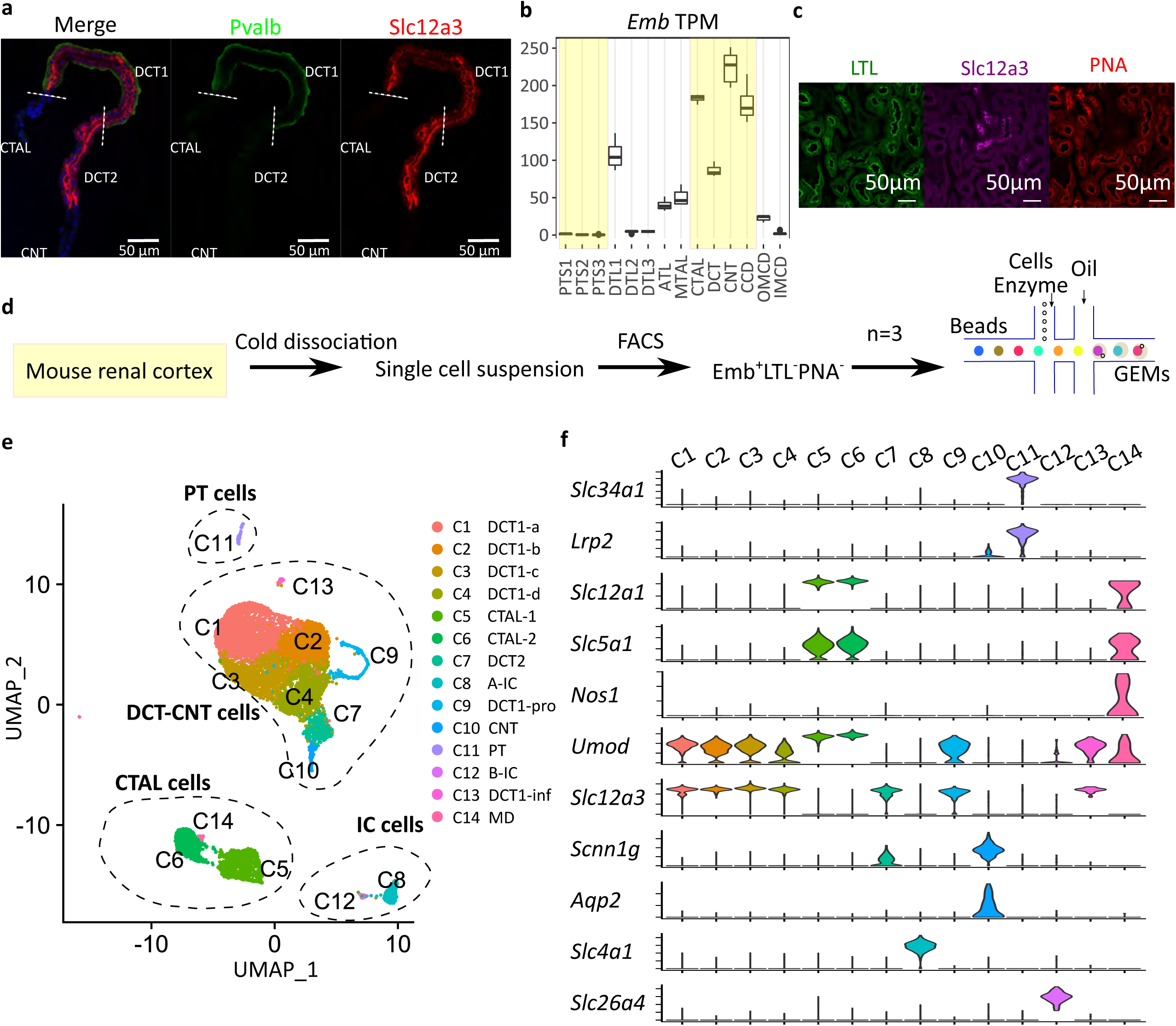
Single-cell RNA-seq of distal convoluted tubule cells. **a** Immunostaining showing the distribution of Slc12a3 (red) and Pvalb (green) in microdissected CTAL to CNT region. Nuclei are stained using DAPI (blue). Scale bars, 50 µm. **b** Distribution pattern of *Embigin* transcript along the mouse renal nephron segments. Embigin was used as a positive surface marker for distal convoluted tubule and thick ascending limb cells, and a negative marker for proximal tubule cells. The tubules in the renal cortex are shown in shaded yellow. **c** Immunostaining showing DCT (Slc12a3 in purple) is negative for LTL (green) and PNA (red). **d** Overview of protocol for the enrichment of distal convoluted tubule cells for single-cell analysis. **e** UMAP projection of cells from all three 10X Chromium datasets. Different clusters are colored and labeled. Four major clusters are highlighted by dashed circles and annotated based on **f. f** Violin plots of renal cell marker genes across all clusters. Cluster information is shown at the top.

Unbiased clustering and UMAP visualization revealed 14 cell clusters corresponding to 5 major cell types, namely PT (C11), CTAL (C5, C6, C14), DCT (C1, C2, C3, C4, C7, C9, C13), IC (C8, C12) and CNT (C10) (Fig. 4d). The transcriptomes of each cell cluster are provided in Supplementary Data 7. The clusters were annotated based on classical renal tubule markers (Fig. 4e) and differentially expressed genes for each cell cluster (Supplementary Data 6). On average, more than 3000 genes/cell were detected in this analysis, and most cells were from DCT+CNT (77.3%), indicating the success of the enrichment process (Supplementary Fig. 3b). Cells sometimes suffer from stress responses during single-cell dissociation characterized by overexpression of immediate early genes, confounding the biological interpretation of scRNA-seq data^42^. As seen in Supplementary Fig. 3c, no evident of expression of the immediate early genes *Fos, Nr4a1*, and *Junb* were found, demonstrating a relative lack of stress artifact in this dataset. Additional analysis of proximal tubule and intercalated cell clusters are provided in Supplementary Fig. 3d. Here we focus first on DCT cell types followed by CTAL subtypes.

### Heterogeneity of DCT1 cells

The scRNA-seq analysis has identified 6 tightly arranged clusters of cells expressing both *Slc12a3* and *Pvalb*, which we identify as DCT1 cells (C1, C2, C3, C4, C9, and C13) (Fig. 5a-b). As shown in Fig. 5c and Supplemental Fig. 4, four of these (C1-C4), expressed the same basic set of genes and are discriminated based on gradients of gene expression, e.g. creatine kinase beta (*Ckb*), decreasing from C1 toward C4; Na^+^/Ca^2+^ exchanger (*Slc8a1*) and calbindin-1 (*Calb1*), decreasing from C2 to C3 (Fig. 5c). The general similarity of the cells in these four clusters leads us to conclude that all of these cells can be classified as a single cell type (‘DCT1 cells’) with subtypes DCT1a-d (Fig. 5c). Two other clusters, C9 and C13, also strongly expressed *Slc12a3* and *Pvalb*, but exhibited superimposed gene signatures that distinguish them from the C1-C4 DCT1 cells. The C9 cells showed marked enrichment of cell cycle and cell proliferation-associated mRNAs (for example, *Stmn1, Mki67, Cenpf*, and *Top2a*) (Fig. 5d and Supplemental Fig. 4). This proliferative profile is consistent with the idea that DCT1 cells are ‘plastic’ in the sense that they can rapidly undergo increases and decreases in cell number and/or volume in response to the delivered load of NaCl from the CTAL^43, 44, 45^. The C13 cells appeared to be DCT1 cells with a superimposed ‘innate immune signature’ with an excess expression of Irf (interferon regulatory factor) transcription factors and many of the genes that they regulate (e.g. MHC class I antigen transcripts and beta-2 microglobulin) (Supplemental Fig. 4). These genes have been associated with damage-associated molecular patterns (DAMPs), triggered by various signals including foreign nucleic acids^46^. These do not appear to represent doublets, e.g. between DCT1 cells and immune cells, because the signatures were not seen with other cell types, e.g. CTAL or proximal tubule. Thus, the expression patterns seen maybe unique to DCT1. In addition to parvalbumin (*Pvalb*), *Lrrc52* (Fig. 5d *right panel*), a subunit of Maxi-K channel^30, 31^, which was selectively found in microdissected DCT in our small-sample RNA-seq, appeared to be expressed in DCT1 cells. Collectively, these data suggest an underlying variation in gene expression among DCT1 cells representing different physiological states of a single cell type.

**Fig. 5.**
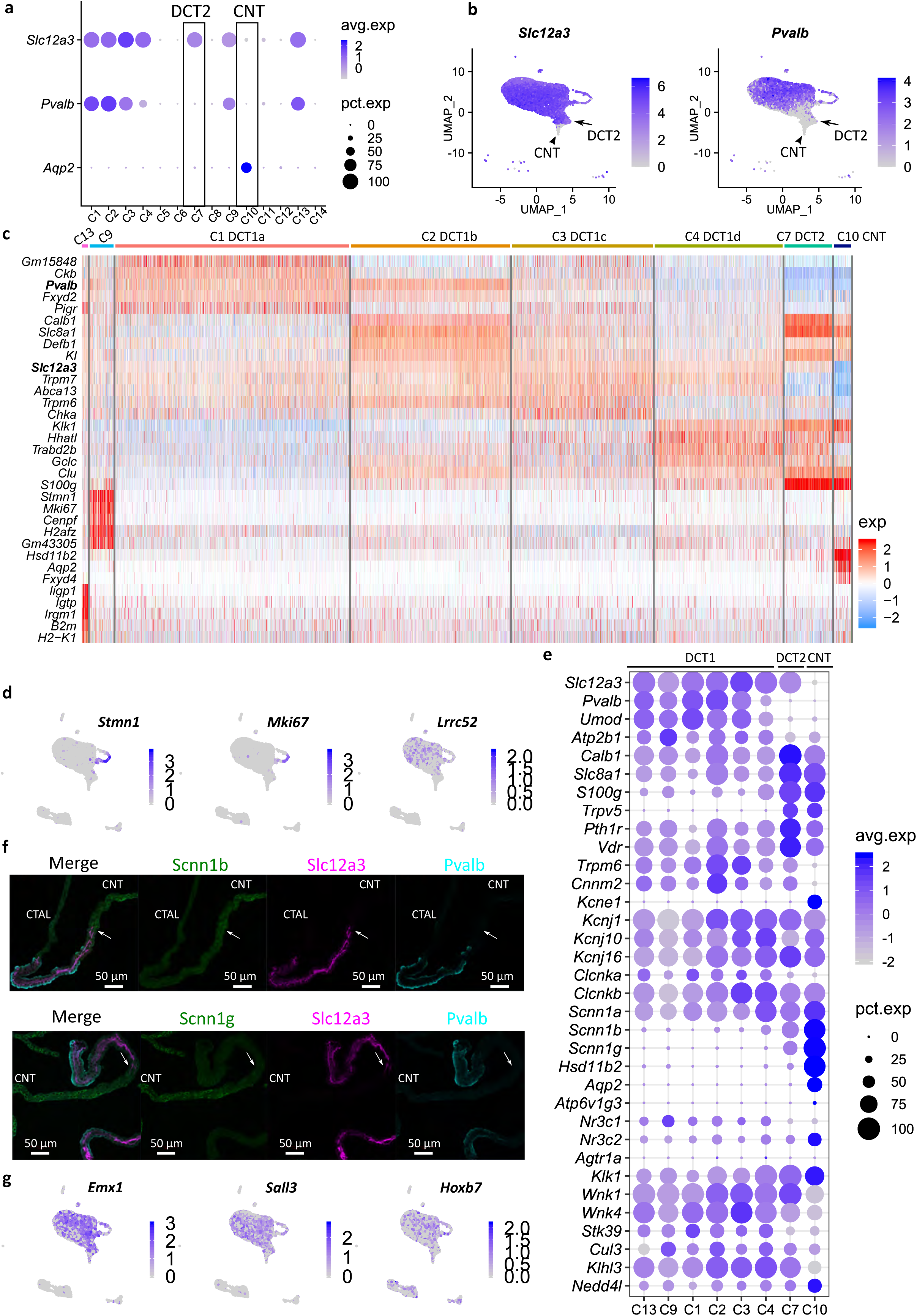
Heterogeneity of distal convoluted tubule cells revealed by single-cell RNA-seq. **a** Dot plot showing the distribution of *Slc12a3, Pvalb*, and *Aqp2* expression across the clusters. Data are normalized and scaled (Z-Score) to examine relative expression across the cell clusters. **b** UMAP projection for *Slc12a3* and *Pvalb* expression. Arrows point to DCT2 cells; Arrowhead points to CNT cells. **c** Heatmap showing the top 5 genes in DCT and CNT cell clusters. Data are scaled (Z-Score). A similar analysis is also provided in Supplementary Fig. 4. **d** UMAP projection for *Stmn1, Mki67, and Lrrc52*. Arrows point to DCT2 cells. **e** Mapping the distribution of major transport pathways in DCT cells. Average expression data are scaled (Z-Score). **f** Immunostaining showing Scnn1b and Scnn1g expression in microdissected tubule from CTAL-DCT-CNT region. Scnn1b and Scnn1g are in green; Slc12a3 is in purple; Pvalb is in cyan, and DNA is in blue. Arrows pointing to regions at point of initial expression of Scnn1b and Scnn1g. Scale bars, 50 µm. **g** UMAP projection for *Emx1, Sall3, and Hoxb7*.

### DCT2 cells and transport pathways

We identify cluster 7 as DCT2 cells. These cells show *Slc12a3* expression without *Pvalb* (Fig. 5a-b), but expressing ENaC (see below). These DCT2 cells can also be distinguished from DCT1 cells in part based on a relative lack of *Umod*^47^ (Fig. 5e). If the numbers of different types of DCT cells profiled are indicative of the relative numbers *in vivo*, the overall ratio of DCT1 to DCT2 cells would be 14.6:1. Thus, DCT2 cells appear to account for a minority of the DCT as a whole. No transcripts were identified that are expressed in DCT2 (i.e. cluster 7) but are not expressed in either DCT1 or CNT (Fig. 5e, Supplemental Fig. 4). That is, no DCT2 specific markers were identified in this study. As the physiology of a particular cell type is determined largely by the genes expressed, this argues against a unique physiological function. Instead, the cells appear to represent hybrid, transitional cells. DCT2 cells strongly express transcripts involved in transcellular calcium reabsorption pathway including two major receptors (*Pth1r* and *Vdr*), an apical calcium channel (*Trpv5*), the intracellular calcium-binding proteins *Calb1* and *S100g*, and the basolateral sodium/calcium exchanger (*Slc8a1*) and plasma membrane calcium-transporting ATPase 1 (*Atp2b1*) (Fig. 5e). In this regard, the DCT2 cells resemble CNT cells and differ from DCT1 in mediating calcium transport^9, 11^. On the other hand, DCT2 resembles DCT1, and not CNT, with regard to the expression of two magnesium transport-related transcripts (*Trpm6* and *Cnnm2*) (Fig. 5e), suggesting that both DCT1 and DCT2 are involved in regulation of the transcellular magnesium reabsorption^48^. Also, supporting this conclusion is that DCT2 is not an independent cell type is the apparent absence of a clear cut DCT2 segment in rabbit kidneys^49^.

Sodium reabsorption in DCT is achieved largely through Slc12a3 (NCC), which is regulated by a signaling pathway involving two Wnk kinases (Wnk1 and Wnk4) and Stk39 (SPAK)^50^. This signaling pathway is believed to be regulated in part by angiotensin II, potassium, and intracellular chloride concentration, and possibly aldosterone^5, 12, 13, 16^. As shown in Fig. 5e, the transcripts coding for major steroid hormone receptors (*Nr3c1* and *Nr3c2*), potassium channels (*Kcnj1, Kcnj10*, and *Kcnj16*), chloride channels (*Clcnka* and *Clcnkb*), and regulatory factors (*Cul3, Klhl3*, and *Nedd4l*) are expressed in both DCT1 and DCT2 cells. Angiotensin II receptor mRNAs (*Agtr1a* and *Agtr1b*), which were undetectable in microdissected DCTs, were also undetectable in DCT1 and DCT2 single cells. DCT2 but not DCT1 cells express *Scnn1b* and *Scnn1g* (Fig. 5e and Fig. 5f). As discussed above with regard to divalent cation transport, the DCT2 transcriptome is not simply the union of the DCT1 and CNT transcriptomes. It lacks expression of the water channel *Aqp2* and *Kcne1*, both of which are strongly expressed in CNT (Fig. 5e). While CNT cells show the highest expression of glucocorticoid metabolizing enzyme *Hsd11b2*, considerably smaller amounts were found in DCT2 cells. Specifically, 96.3% of CNT cells express *Hsd11b2* at an average expression at 33.2 in log scale. However, only 23.3% of DCT2 cells had detectable levels with a much lower average value (0.48 in log scale). DCT1 cells (C1-4) had barely measurable levels in less than 6% of cells (Fig. 5e).

An alternative to a purely physiological role for DCT2 cells is the possibility that the role of these cells is to join the nephron and collecting duct which are derived from different developmental progenitors^51^. Some clues to a possible such role could be derived from patterns of expression of homeobox genes that are generally involved in tissue segmentation during development. One homeobox gene (*Emx1*) was found to be highly selectively expressed in DCT in our small-sample RNA-seq profiling, while the single-cell analysis did not show any selectivity in DCT2 versus DCT1 cells (Fig. 5g, *left panel*). Emx1 has previously been identified in zebrafish as an important regulator of distal nephron development^34^. *Sall3*, which was identified as DCT-selective in the small-sample RNA-seq studies, is also not selective for DCT2 versus DCT1 (Fig. 5g, *mid panel*). The homeobox gene most associated with ureteric bud differentiation is *Hoxb7*^52^, which is approximately equally expressed in all DCT cell subtypes, CNT, and intercalated cells (Fig. 5g, *right panel*).

### Mosaic cell type pattern in cortical thick ascending limb (CTAL)

Single-cell RNA-seq analysis identified three distinct CTAL (*Slc12a1*^+^) cell subtypes (C5, C6, and C14) (Fig. 6a). One of these (C14) expressed genes characteristic of macula densa cells, namely *Nos1*^*53, 54*^ and *Avpr1a*^55^ (Fig. 6b and Supplementary Fig. 5a). We found that *Pappa2* was also strongly expressed in MD cells (Fig. 6b). This gene has been previously linked to salt-sensitive hypertension in Dahl S rats^56^. The full MD transcriptome is provided in Supplemental Data 7 (See *Method*). Strikingly, these cells expressed multiple neuronal genes aside from *Nos1*, including several involved with axon guidance and synaptic adhesion (*Robo2, Ncam1*, and *Auts*). This suggest the characterization of MD cells as ‘neuroepithelial cells’. We speculate that the axon-guidance proteins could be part of the homing mechanism that localizes MD cells to the vascular pole of the glomerulus. Interestingly, MD cells selectively express *Bmp3* (Supplementary Fig. 5b), a secreted protein that could be involved with communication with other cell types including DCT cells if secreted apically or renin-secreting juxtaglomerular cells (JG cells) if secreted basolaterally. In addition, there are multiple transcripts that are strongly expressed in the CTAL but are very low in macula densa including uromodulin (*Umod*)^57^ and Egf (*Egf*)^32^ (Fig. 6c and Supplementary Fig. 5a).

**Fig. 6.**
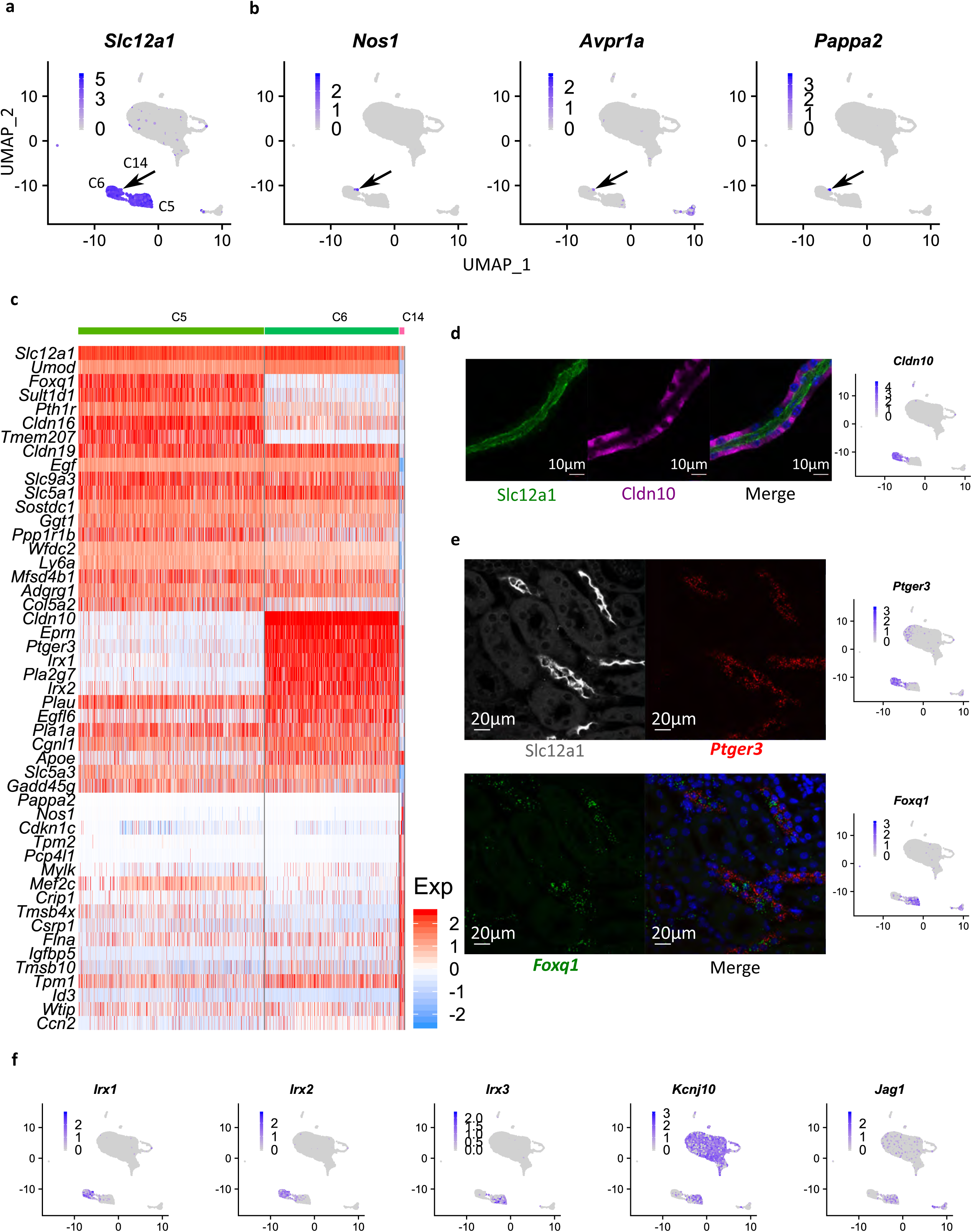
Single-cell analysis of cortical thick ascending limb cells. **a** UMAP visualization of Slc12a1. **b** UMAP plot for *Nos1, Avpr1a, and Pappa2*. **c** Heatmap showing the top 20 marker genes in C5, C6, and C14. Each column indicates a cell. Colors indicate the expression level (Z-Score). **d** Immunostaining showing Slc12a1 (green) and Cldn10 (purple) on a microdissected mouse TAL. Nuclei are stained with DAPI (blue). The feature plot for *Cldn10* is shown on the right. Scale bar, 10 µm. **e** Immunostaining coupled with in situ hybridization (RNAscope) showing Slc12a1 (white), *Ptger3* (red), and *Foxq1* (green) on mouse kidney section. Nuclei are stained with DAPI (blue). Feature plots are shown on the right. Scale bar, 20 µm. **f** UMAP visualization of *Irx1, Irx2, Irx3, Kcnj10*, and *Jag1*.

The other two CTAL clusters were distinguished by *Cldn10* and *Ptger3* in one (C6) and *Cldn16* and *Foxq1* in the other (C5). As shown in Fig. 6d, some CTAL cells lack Cldn10 as reported before^58^. Multiplex localization of Slc12a1 (immunofluorescence) with *Foxq1* and *Ptger3* mRNA (in situ hybridization with RNAScope) in mouse kidney sections highlighted the mutually exclusive pattern of *Foxq1* and *Ptger3* in TAL (Fig. 6e). In addition, these two CTAL cell types were distinguished by alternative expression of Iroquois homeobox transcription factors, with *Irx1* and *Irx2* in the *Cldn10*^+^ CTAL cells (C6) and *Irx3* in the *Foxq1*^+^ CTAL cells (C5) (Fig. 6f). The subsequent analysis showed that *Jag1* (ligand for Notch) was predominantly expressed in C5 (Fig. 6e), while *Hes1* expression was greatest in C6 (Supplementary Fig. 5c), suggesting a potential binary cell fate between these two CTAL cell types regulated by Notch signaling^59^. Interestingly, C5 also strongly expressed *Kcnj10* but not *Kcnj16*, whereas both are expressed in other distal cell types where they co-assemble into heterodimeric channels^60^ (Fig. 6f).

## Discussion

A complete understanding of the cellular composition and transcriptional profiles of the cells in DCT has been lacking. Here, we combined cell enrichment and single-cell RNA-seq to investigate the transcriptome landscape of *Slc12a3*^+^-DCT cells (https://esbl.nhlbi.nih.gov/MRECA/DCT/). Our data demonstrate that DCT cells are subdivided into *Slc12a3*^+^/*Pvalb*^+^ DCT1 cells and *Slc12a3*^+^/*Pvalb*^-^ DCT2 cells. We observed that DCT1 cells are heterogeneous. This heterogeneity appears to be associated with variable expression of *Slc8a1, Calb1*, and *Ckb* among other mRNAs. The relevance of such topology of gene expression to the spatial arrangement of DCT1 cells, as well as DCT function, remains for further investigation. How can the heterogeneity of DCT1 cells or the hybrid property of DCT2 cells (mentioned below) be explained? Since the DCT and downstream collecting duct system are derived from two separate progenitors, one possible explanation for the heterogeneity could be reciprocal mutual inductions between nephron progenitors and ureteric bud during kidney development^51^. Among DCT1 cells, we also identified a small population of cells that showed marked enrichment of cell cycle and cell proliferation-associated mRNAs (e.g. *Pcna, Mki67, Cdk1*, and *Top2a*). These cells are rare but unlikely to be artificial doublets as these proliferation markers are not seen in other epithelial cells (PT, CTAL, and ICs) studied in this experiment. This finding suggests that certain DCT1 cells might possess a proliferative potential, even in normal kidneys under normal physiological conditions. Consistent this conclusion is that the DCT is known to undergo extensive remodeling under different stimuli, especially changes in luminal NaCl delivery^43, 44, 45^. A future direction would be to assess how these cells adapt to changes in luminal NaCl load and whether this proliferative-type DCT1 cell undergoes changes in abundance in response to such stimuli.

Identification of unique DCT2 markers could be the basis of future methods for isolation, identification, and targeting of DCT2. However, no unique transcripts were found in these cells. Thus, DCT2 cells are more likely to be hybrid cells between DCT1 and CNT cells. Our data suggest that similar to DCT1 cells, DCT2 cells are involved in sodium (*Slc12a3 and* ENaC) and magnesium reabsorption (*Trmp6* and *Cnnm2*). Our study revealed that DCT2 cells had detectable but much lower levels of *Hsd11b2* mRNA than CNT. However, the experiments don’t reveal protein levels or enzyme activity in DCT2 cells. In a recent study using protein mass spectrometry in microdissected tubules of rat, we found readily measurable Hsd11b2 protein levels in DCT (chiefly DCT1)^61^. Thus we cannot rule out the possibility that there is enough Hsd11b2 in DCT1/DCT2 to metabolize glucocorticoids. Furthermore, Fig. 5e shows significant mineralocorticoid receptor (*Nr3c2*) expression in all DCT clusters. In addition to their ENaC expression, the DCT2 cells resemble CNT cells by strongly expressing transcripts coding for apical Ca^2+^ entry protein (*Trpv5*) and basolateral Ca^2+^ extrusion proteins (*Atp2b1* and *Slc8a1*). Thus, our study supports the observation that DCT2 cells have an important role in transcellular calcium reabsorption^9, 11^.

We found three types of CTAL cells, including a small cell cluster with a gene expression pattern consistent with macula densa (MD) cells. These cells express the classical MD cell marker, neuronal nitric oxide synthase (Nos1)^53, 54^, as well as a number of neuronal genes making these cells as ‘neuroepithelial’ in character. Identification of transcripts coding for secreted proteins like *Pappa2* and *Bmp3* may shed light on the ways that MD cells communicate with neighboring cell types. The other two CTAL cell types were more abundant. Previous studies found two morphologically distinguishable cell types in rat TAL: rough-surfaced cells (RSC) with well-developed microvilli and smooth-surfaced cells (SCC)^62^. These cells differ in apical/basolateral K^+^ conductance^63, 64, 65^ and subcellular localization of *Slc12a1* (NKCC2)^66^ and *Egf*^67^. A striking observation from the present study is the identification of two CTAL subtypes based on gene expression that might correspond to the morphologically identified cell types, namely cells corresponding to the C5 cluster (*Foxq1*^+^*Cldn10*^-^*Ptger3*^-^) and the C6 cluster (*Foxq1*^-^ *Cldn10*^+^*Ptger3*^+^). C5 was strongly enriched in *Kcnj10* but not *Kcnj16*. Whether Kcnj10 alone mediates the basolateral K^+^ conductance in this CTAL subtype remains for further investigation^68^. Our data further showed *Jag1* was expressed in C5 while absent in C6. Such a mosaic cell pattern mimics collecting duct epithelium (i.e. PC and IC), where cell specification is regulated by the Notch signaling pathway^69^. Whether the Notch signaling pathway regulates the fate of these two cell types in TAL remains unknown. Also, the differential expression of *Irx1*/*Irx2* versus *Irx3* in these two cell types could play roles in their differentiation and physiological function.

Finally, we provide a comprehensive gene expression atlas for 14 mouse renal tubules by small-sample RNA-seq, available at https://esbl.nhlbi.nih.gov/MRECA/Nephron/. Given the current scope of the manuscript, we only focused on certain aspects of these data. They are likely to be useful for a wide number of future applications. Another important aspect of these data is that it adds information on alternative splicing differences along the renal tubule, adding to other recent evidence^70^. We believe these data together with previous RNA-seq^17^ and more recent proteomic data^61^ from rat tubules will enhance our understanding of renal function.

In summary, systematic analyses of cellular composition and gene expression provide insights into kidney functions and diseases. We provide our data in a comprehensive website and expect that this study will be an essential asset for future kidney studies. Since technological advances enable us to profile different properties of tissues or individual cells, such as the genome, transcriptome, epigenome, and proteome, or even multi-omes^71^, we expect these approaches will open avenues for further kidney research.

## Methods

### Adult mouse kidney cell isolation and single-cell RNA-seq

This protocol was performed as described previously^24^. The kidneys were perfused directly via the left ventricle with perfusion buffer (5mM HEPES, 120mM NaCl, 5mM KCl, 2mM CaCl_2_, 2.5mM Na_2_HPO_4_, 1.2mM MgSO_4_, 5.5mM glucose, 5mM Na acetate in H2O at pH 7.4) to remove blood cells, followed by perfusion with the same perfusion buffer supplemented with 1mg/ml Collagenase B (Sigma 11088831001). Kidneys were immediately removed and placed in ice-cold PBS. The cortical region of kidney tissue was then dissected and minced on ice, followed by dissociation in 20ml cold perfusion buffer consisting of collagenase B (2.5 mg ml^-1^), Dispase II (1.2U ml^-1^, Roche), 7.5 mg/ml B. licheniformis protease (Sigma # P5380) and DNase I (125 Kunitz ml^-1^, Worthington, #LS002058) for 25-30 min at 10°C with frequent agitation at every 5min. In the last 10min before the end of digestion, the suspension was passed through a 20 gauge needle. Digestion enzymes were neutralized by adding 20ml 20% FBS and the suspensions were then centrifuged at 250g at 4°C for 3min. The pelleted cells were resuspended briefly in 5ml lysis buffer (Quality Biological,118-156-101) for 30s to remove red blood cells followed by dilution in 45ml PBS. After centrifugation at 250g for 3min, the pellet was resuspended and washed again with 50ml PBS. The final cell pellet was resuspended in 30ml FACS buffer (perfusion buffer with 0.05% BSA) and then passed sequentially through 100 µm, 70 µm and 40 µm cell strainers (VWR) to obtain a suspension of isolated cells (single-cell suspension). The single-cell suspension was pelleted at 250g for 3min and resuspended in 1ml FACS buffer. 120µl FcR blocking reagent (Miltenyi Biotec, 130-092-575) was added to 1ml single-cell suspension together with 1:500 Hoechst 33342 (Invitrogen, #H3570), 1:200 LTL-FITC (Vector Laboratories, FL-1321-2), 1:500 PNA-Cy5 (Vector Laboratories, CL-1075-1), 1;200 CD45 (BioLegend, 103149), 1:10 Emb-PE (Miltenyi Biotec, 130-107-948). The staining was done in the dark at 4°C with agitation. After 30min staining, the cells were washed twice with FACS buffer and resuspended in 2ml FACS buffer before sorting. 7-AAD (BD, #559925) at 1:10 was added to the cells before FACS. The cells were sorted in 700µl FACS buffer with a low flow rate and pressure to ensure high viability. 30-40K cells were collected and centrifuged at 250g for 3min in a swinging bucket at 4°C. The top supernatant was carefully removed and 30 µl volume was left to resuspend the cell pellet. 10µl cells were mixed with 10µl Trypan Blue (Lonza, #17-942E) and then loaded on Countess II Automated Cell Counter (Invitrogen) for counting and viability test. Cells at 700-1200 cells/µl were directly loaded to 10X chips for Gel Bead-In Emulsions (GEM) generation.

### Single-cell RNA-seq data analysis

Lysed or dead cells in droplet-based single-cell RNA-seq assays release ambient RNA or cell-free RNA, resulting in a low Fraction Reads in Cells (FRC), i.e. reads are not 100% associated with a valid cell barcode in GEM. Such background levels of contamination vary between batches and cell types. SoupX (https://doi.org/10.1101/303727) estimates such ambient RNA by pooling of droplets with low numbers of UMI (10 UMI or less). As the majority of generated GEMs contain no cells, this provides means to estimate the background noise. The bimodal distribution of transcripts guides the contamination fraction calculation for each cell. The counts are then adjusted for downstream analysis.

10X Chromium raw sequencing data were first processed with cellranger (v 3.0.2). SoupX (V 0.3.1) was then applied for background correction. *Kap* and *Slc12a1* were identified and used for expression profile correction. 3 batches were independently estimated, corrected, and merged in R with Seurat (V 3.0) package. Initially, cells with less than 200 features (genes), and features expressed in less than 10 cells were discarded. Cells were further filtered based on :nFeature_RNA > 1000 & nFeature_RNA < 5800 & percent.mt < 35 & percent.mt >5 & nCount_RNA <30000. A small cluster of cells showing extremely low mitochondria reads and UMI were removed. The resulting Seurat object contains 17655 features across 9099 samples. To correct batch effects for 3 independent preparations, the integration workflow was applied to the combined object. The combined object was split, normalized, and the top 2000 highly variable features were identified. Anchors were identified using the FindIntegrationAnchors function, which used the first 30 Canonical Correlation Analysis (CCA) dimensions for searching. Datasets were then integrated based on the anchors found before.

Standard Seurat analysis was then applied to the integrated data. Data were scaled and mitochondria variation was regressed out. The first 15 principal components were chosen for downstream clustering after linear dimensional reduction (RunPCA). Cell clusters were determined by FindNeighbors and FindClusters. Uniform Manifold Approximation and Projection (UMAP) was used throughout the paper to visualize the single-cell data.

Differentially expressed genes in each cluster were calculated by FindAllMarkers function. For better visualization, sometimes subset function was used to extract certain clusters. Other R packages including ggplot2 and dplyr were used for final figure preparation.

### Mouse renal tubule microdissection

6-to 8-week-old male C57BL/6 mice (Taconic) were used for renal tubule microdissection. All animal work in this study was conducted in accordance with NIH animal protocol H-0047R4. Mice were sacrificed via cervical dislocation. The kidneys were perfused via the left ventricle with ice-cold DPBS (Thermo Scientific) to remove blood cells, followed by reperfusion with the dissection buffer (5mM HEPES, 120mM NaCl, 5mM KCl, 2mM CaCl_2_, 2.5mM Na_2_HPO_4_, 1.2mM MgSO_4_, 5.5mM glucose, 5mM Na acetate, pH 7.4) with 1 mg/ml Collagenase B (Roche). Kidneys were harvested and thin tissue slices along the cortical-medullary axis were obtained and proceeded to digestion. The dissociation was carried out in dissection buffer containing collagenase B (1-2 mg/ml) and hyaluronidase (1.2-2 mg/ml, Worthington Biochemical) at 37 °C with frequent agitation for 30 min. The digestion was monitored until the optimal microdissectable condition was reached. Most renal tubules were readily microdissectable under this condition except the inner medullary collecting ducts (IMCDs), which required a higher enzyme concentration (2 mg/ml) and a longer digestion time (up to 90 min at 37 °C). The microdissection was carried out under a Wild M8 dissection stereomicroscope equipped with on-stage cooling. The renal tubules were washed in dishes containing ice-cold DPBS by pipetting to remove contaminant before RNA extraction using a Direct-zol RNA MicroPrep kit (Zymo Research, Irvine, CA). 4-8 tubules were collected for each sample.

### Small-sample RNA-seq

These steps were performed as previously reported^19^. RNA was eluted in 10 µl water and immediately proceeded to cDNA generation using the SMART-Seq V4 Ultra Low RNA Kit (Takara Bio USA, Mountain View, CA). 1 ng of the resulting cDNA from 14 PCR cycles was “tagmented” and barcoded by using a Nextera XT DNA Sample Preparation Kit (Illumina). The final libraries were purified by AmPure XP magnetic beads (Beckman Coulter, Indianapolis, IN), and quantified using a Qubit 2.0 Fluorometer (Thermo Fisher Scientific, Waltham, MA). An equal amount of index libraries were pooled and sequenced (paired-end 50 bp) on an Illumina Hiseq 3000 platform.

RNA-seq reads were aligned by STAR(2.5.4a)^72^ to the mouse Ensembl genome (release 94) with the corresponding annotation file (release 94). The unique genomic alignment was processed for alignment visualization on the University of California, Santa Cruz, Genome Browser, or Jbrowse^73^. Transcript per million (TPM) and expected counts were obtained using RSEM(1.3.0)^74^. Samples with fewer than 75% uniquely mapped reads and less than 10 million uniquely mapped reads were excluded for downstream analysis. Unless otherwise specified, the computational analyses were performed on the National Institutes of Health Biowulf High-Performance Computing platform.

### Differential expression analysis and hierarchical clustering

Differentially expressed genes (DEGs) were identified using R package edgeR^75^. Row expected counts (RSEM output) were as an input for this analysis. Each renal tubule segment was compared against the rest of the segments. Genes from these comparisons with max TPM >= 25, absolute logFC >=2, and FDR < 0.05 were merged. A total of 3709 non-redundant genes were included in the following analysis. Hierarchical clustering analysis was done in R using the Pearson correlation. TPM values were used in the log_2_(TPM+1) scale. Distance between samples or between genes is defined as as.dist(1-correlation). Heatmaps were generated using the heatmaply package.

### Isoform analysis

Alternative splicing analysis was performed with MAJIQ 2.1^41^. To facilitate the local splicing variations (LSVs) identification process, RNA-seq data were remapped using STAR in a 2-pass mode^72^ before MAJIQ analysis. Exon’s percent spliced in (PSI, or Ψ) was estimated for each renal segment with replicates. Similar to gene differential analysis described above, the individual renal segment was compared to other segments to estimate the change in each splicing junction’s relative inclusion level, termed delta PSI (ΔΨ). High-confidence differential splicing variants were selected based on P(ΔΨ > 0.2) > 0.95. All non-redundant differentially spliced LSVs along the renal tubule were included in the final list. Splicing variants were visualized using Voila^41^. Representative LSVs from each renal segments were further analyzed with MAJIQ-SPEL^41^ on Galaxy server https://galaxy.biociphers.org/galaxy/root?tool_id=majiq_spel. The alignments were visualized on the University of California, Santa Cruz, Genome Browser, and tracks were downloaded for figure preparation.

### Immunohistochemistry and RNAscope in situ hybridization

The mouse kidney tissue was prepared as described previously^19^. Mice were cervically dislocated and perfused with ice-cold DPBS followed by 4% PFA in DPBS. Kidneys were then fixed for two hours in 4% PFA before transferring to 20% sucrose at 4°C overnight. The blocks were further processed in NHLBI Pathology Facility. 6-µm-thick sections were cut. Frozen sections were thawed at room temperature for 10-20 minutes and rehydrated in PBS for 10 minutes. After blocking for 30 min with 1% BSA and 0.2% gelatin, primary antibodies were applied overnight at 4°C. Sections were washed for 3 × 5 min in PBS. The secondary antibody incubation was carried out for 1 hour at room temperature.

RNAscope was performed on fixed frozen tissue sections according to RNAscope Multiplex Fluorescent Reagent Kit v2 user manual (document #323100-USM). TSA^®^ Plus fluorophores at the concentration of 1:1500 were used to develop signals. Once the signals were developed for each channel, standard immunohistochemistry protocol was followed for antibody staining. The confocal images were taken in the NHLBI Light Microscopy Core.

### Immunohistochemistry antibodies and RNAscope probes

Immunolabeling reagent includes several antibodies generated in the Knepper lab include rabbit anti-Slc12a3 (4375), chicken anti-Slc12a1 (C20), chicken anti-Scnn1g (LC38), and chicken anti-Scnn1b (LC40). Commercial antibodies were Pvalb (Swant, GP72) and Cldn10 (Thermo Fisher Scientific, Catalog # 38-8400). Probes used in this study are Foxq1 (Advanced Cell Diagnostics, Cat No. 504801) and Ptger3 (Advanced Cell Diagnostics, Cat No. 501831-C3). Fluorescently labeled lectins were LTL (Vector Laboratories, FL-1321) and PNA (Vector Laboratories, CL-1073 or CL-1075).

### Websites

Websites were built around the Shiny package in R. For https://esbl.nhlbi.nih.gov/MRECA/Nephron/, data tables were displayed using the DT package. The plotly, ggplot2, and heatmaply were used to create the interactive heatmaps or feature plots. RSQLite, dbplyr, and pool packages were employed to write and fetch data from GTEx V8 RNA-seq database. https://esbl.nhlbi.nih.gov/MRECA/DCT/ was developed in Shiny to directly visualize single-cell RNA-seq data. Interactive UMAP and Dot plot were create using Seurat and modified by ggplot.

### Data availability

Sequencing data including RNA-seq and single-cell RNA-seq reported in this paper have been deposited in the Gene Expression Omnibus (GEO, https://www.ncbi.nlm.nih.gov/geo/query/acc.cgi?acc=GSE150338, Temporary reviewer security token: gjkrumgwbbitdyh). The source data underlying Figs. 2a,d, 3a-b, 4a and Supplementary Figs. 1a-b,d-e are provided as a Source Data file.

## Acknowledgments

The work was primarily funded by the Division of Intramural Research, National Heart, Lung, and Blood Institute (project ZIA-HL001285 and ZIA-HL006129, M.A.K.). Next-generation sequencing was done in the NHLBI DNA Sequencing Core Facility (Yuesheng Li, Director). Confocal images were taken in the NHLBI Confocal Microscopy Core Facility (Christian Combs, Director). Cell sorting was performed in the NHLBI Flow Cytometry Core (J. Philip McCoy, Director). Cell viability assay and counting were done in NHLBI induced Pluripotent Stem Cells Core (Jizhong Zhou, Director). Tissue sections were prepared in the NHBLI Pathology Facility (Zu-Xi Yu, Director).

## Author contributions

L.C. and M.A.K. designed research, analyzed data, and made figures; L.C. performed experiments and designed websites; L.C., C.-L.C., and M.A.K. contributed reagents, discussed the results, and drafted the manuscript. The authors declare that there is no conflict of interest.

