## Supplementary figures and images for "Targeted single-cell RNA-seq identifies minority cell types of kidney distal nephron that regulate blood pressure and calcium balance"

### Supplementary Fig. 1

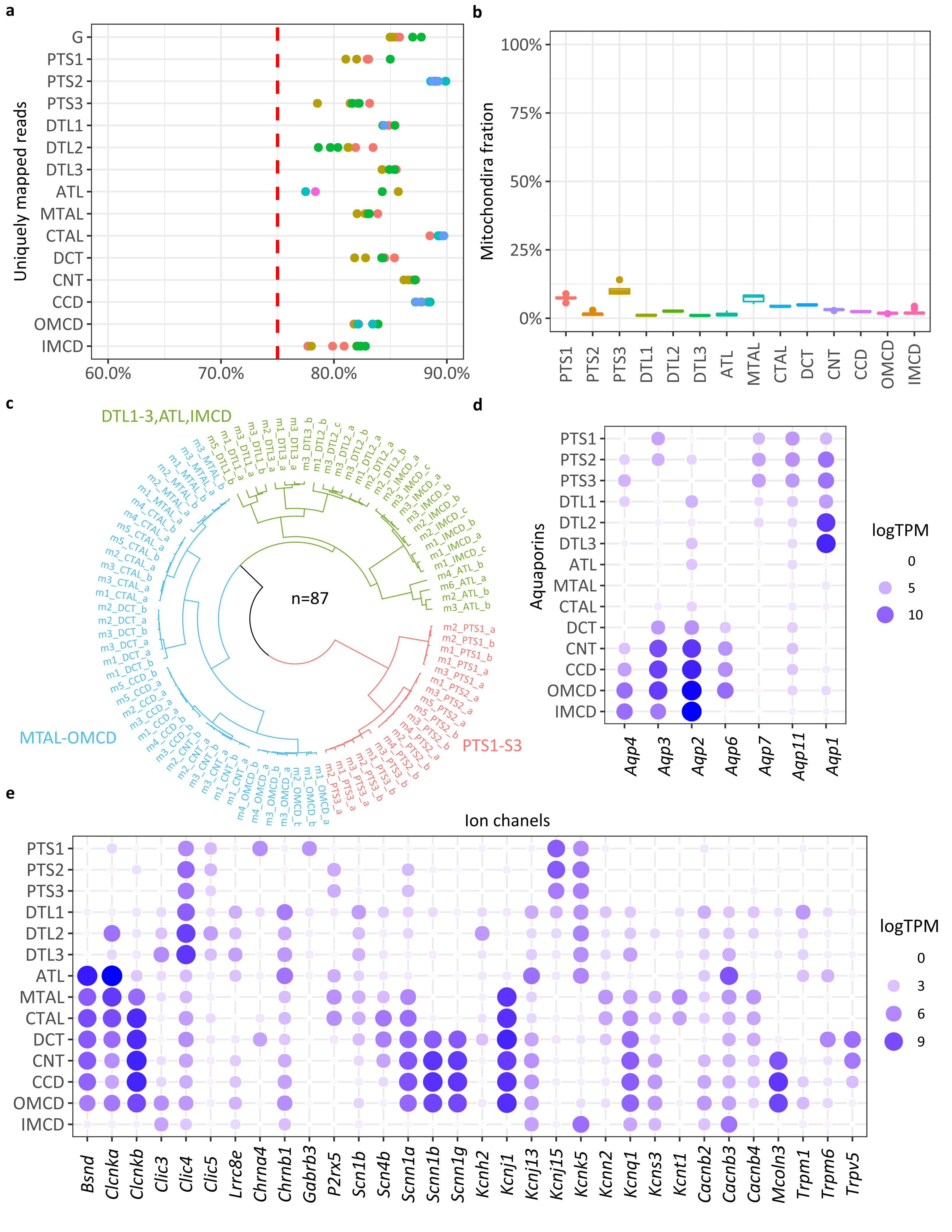

### Supplementary Fig. 2

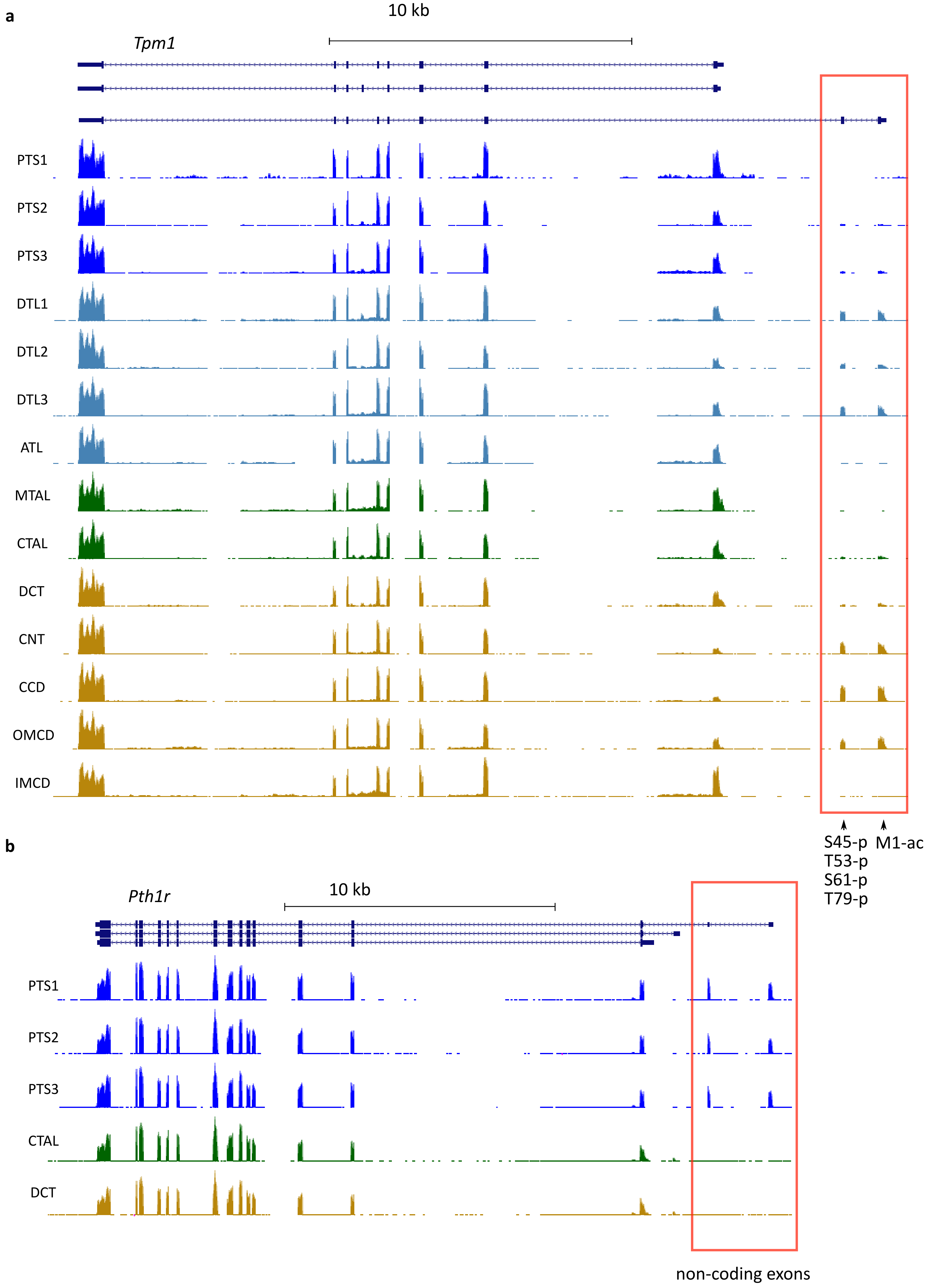

### Supplementary Fig. 3

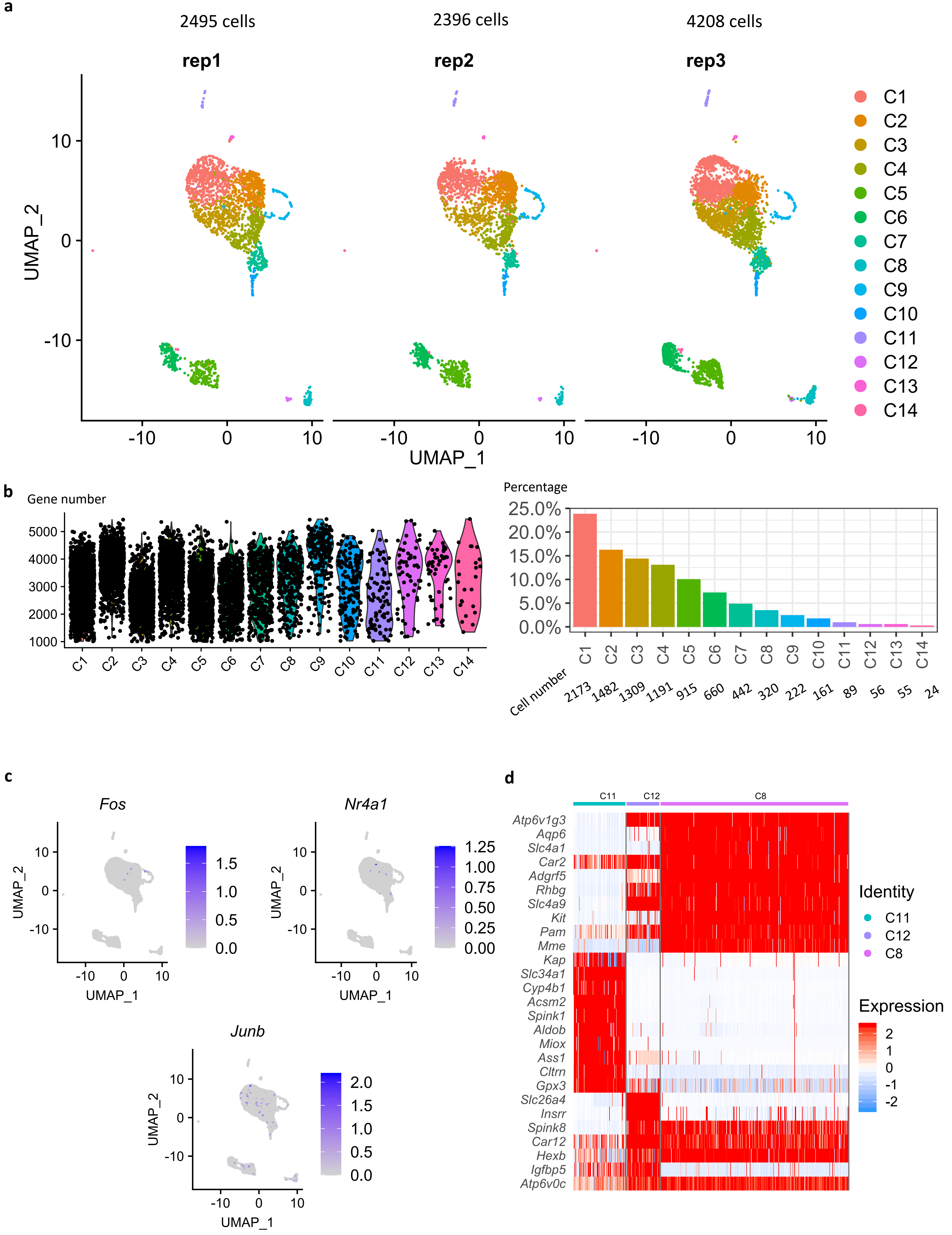

### Supplementary Fig. 4

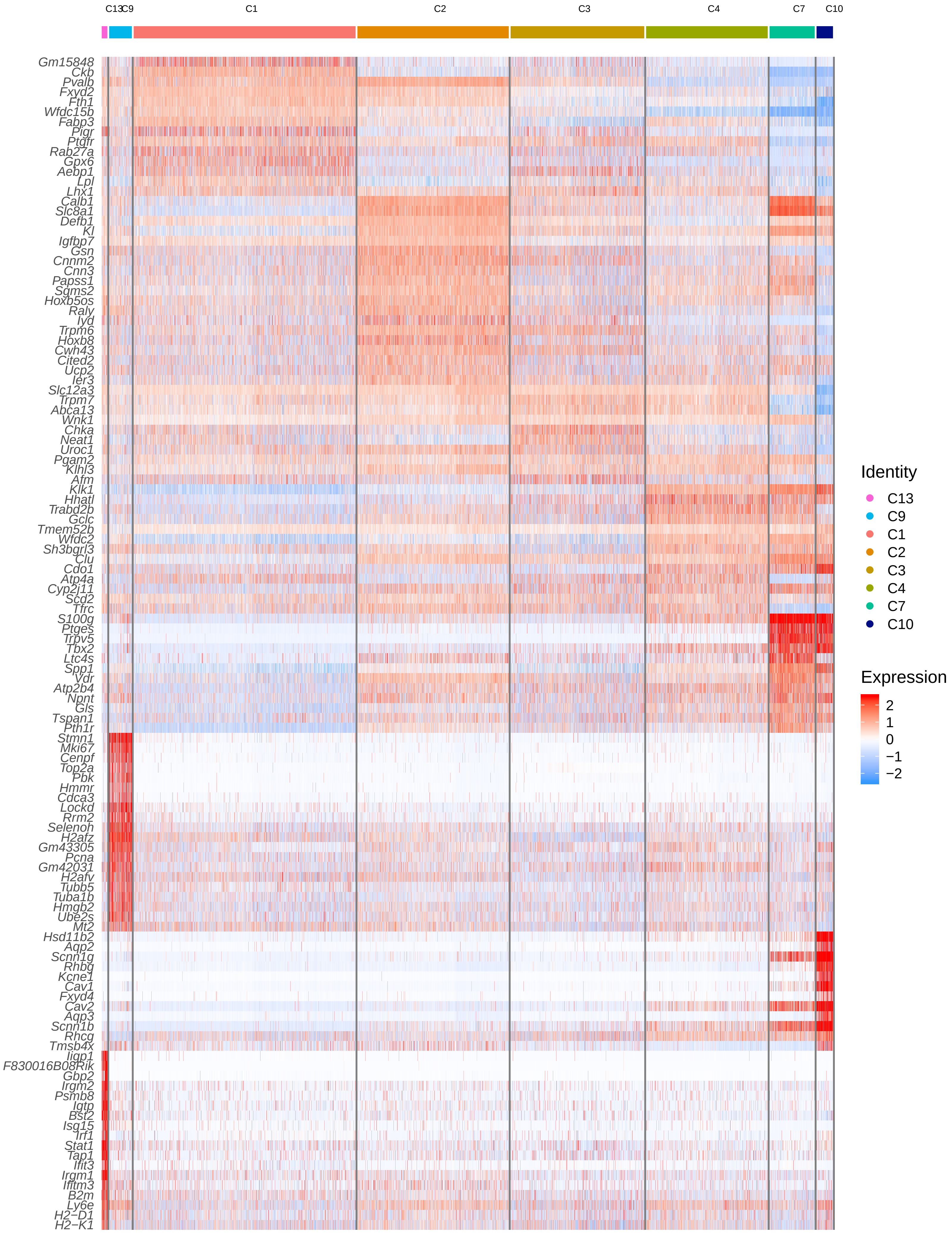

### Supplementary Fig. 5

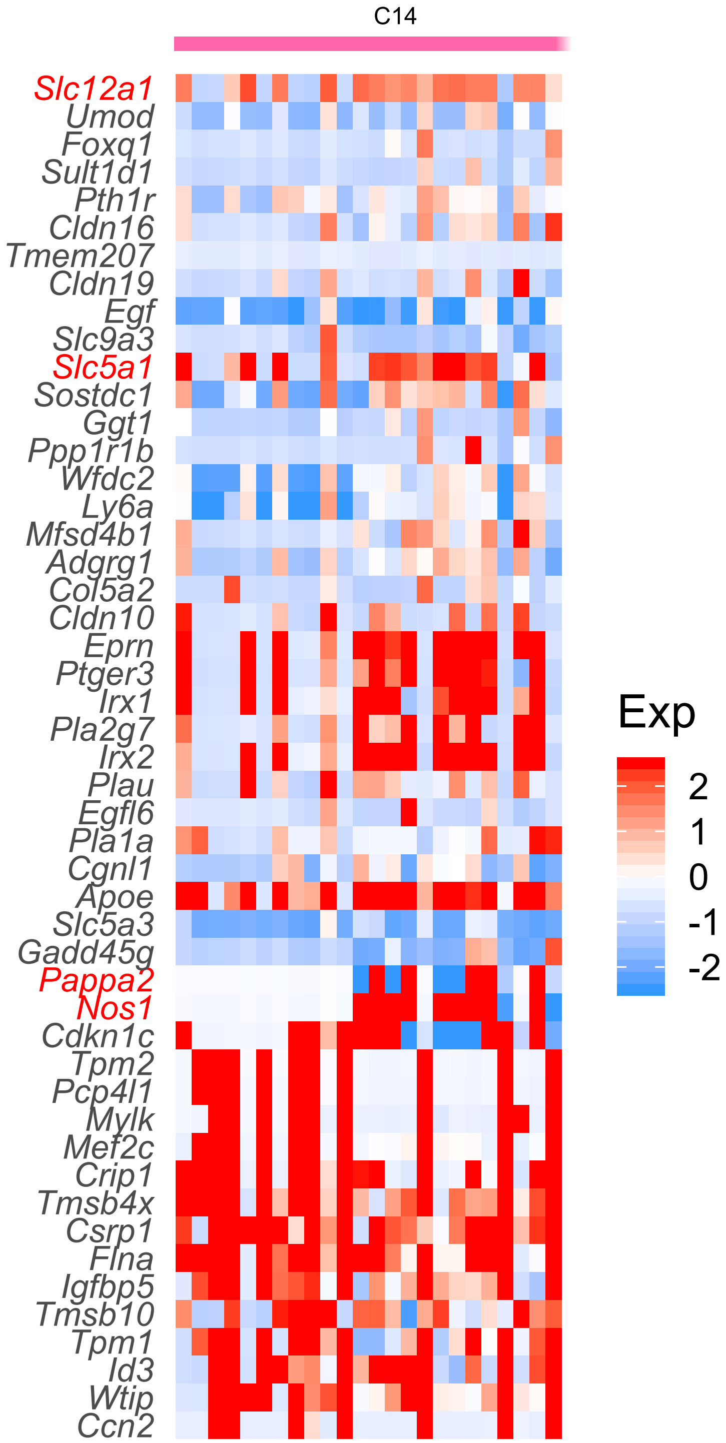

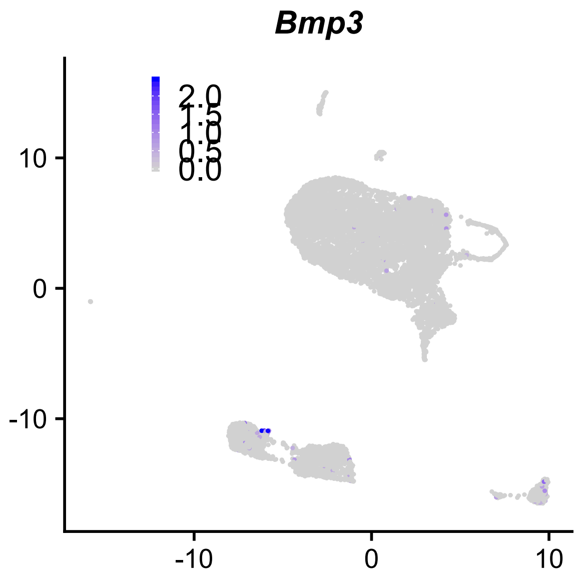

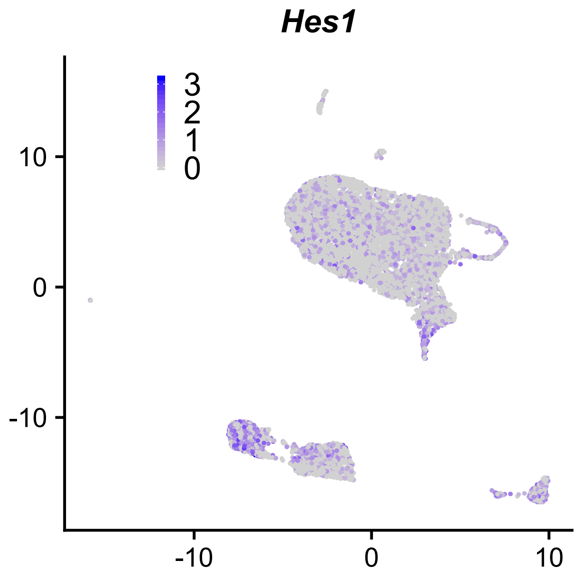
